# Socially dominant male mice in social hierarchies identified via automated RFID tracking exhibit elevated activity levels and circulating markers of higher metabolic demand

**DOI:** 10.64898/2026.08.26.747333

**Authors:** SO Seese, TM Milewski, M Fusillo, JP Curley

## Abstract

Dominance hierarchies are a fundamental aspect of social organization, enabling animals to minimize aggression and optimize access to resources. Previous studies have highlighted the energetic and physiological demands of dominant status, as well as the behavioral flexibility required of subordinates to navigate these hierarchies. Despite advancements in automated behavior tracking, limitations persist in tracking fine-scale, real-time interactions within complex social environments. Here, we developed and validated a novel RFID-based system to continuously monitor dominance hierarchies in group-housed male mice over 10 days. This system enabled unbiased behavioral inference across light phases and revealed spatial and temporal patterns of dominance behavior undetectable through traditional live-scored methods. Automated tracking accurately identified alpha individuals and consistently inferred linear hierarchies across cohorts, with greater precision for higher-ranked individuals. Behavioral metrics, such as transition frequencies and proximity to food zones, were consistent with dominance driven activity. Hormonal analyses revealed that higher-ranked mice exhibited increased leptin and peptide YY, consistent with heightened activity and satiety signaling, while lower C-peptide levels reflected greater metabolic demands of dominance. Furthermore, dominance rank was associated with differences in light–dark activity, which were in turn related to circulating hormone profiles. This study demonstrates the utility of automated RFID tracking in capturing dominance hierarchies with temporal and spatial granularity, while revealing links between social rank, metabolic regulation, and activity patterns advancing our understanding of social behavior dynamics.

## Introduction

A foundational aspect of social behavior across a wide range of species is the establishment and maintenance of dominance hierarchies (Chase, 1982). These hierarchies are among the most thoroughly studied forms of social organization, emerging through competition for access to limited resources such as food, territory, or reproductive opportunities (Chase & Seitz, 2011). Within these social structures, it is critical that individuals assess their own rank and monitor the relative status of others to adaptively regulate behavior, reduce unnecessary conflict, and maintain group stability (Drews, 1993). This regulation depends on the integration of external cues, such as recognition of others and memory of social context, with internal physiological states to guide status-appropriate behaviors (P. Chen & Hong, 2018; Milewski et al., 2022; Taborsky & Oliveira, 2012). These recognition-driven and context-sensitive social interactions are not only essential for conflict mitigation but are also key to survival and reproductive fitness. Once hierarchies are established, individuals must quickly recognize rank relationships and modulate their own behavioral strategies, becoming more dominant or more submissive, depending on their relative position within the group (Anderson et al., 2016; Chase, 1982; P. Chen & Hong, 2018; Insel & Fernald, 2004; O’Connell & Hofmann, 2011).

Dominant individuals, particularly those at the top of the hierarchy, often incur substantial energetic costs in acquiring and maintaining their status. In many species, dominant males must perform energetically costly behaviors including aggressive encounters, territory patrolling, and frequent dominance signaling (Milewski et al., 2022). Similar energy-intensive patterns occur across species. In chimpanzees, high-ranking males excrete lower levels of C-peptide, a biomarker associated with insulin secretion, indicating negative energy balance and high metabolic output (Thompson et al., 2009). In rhesus macaques, dominant males show reduced C-peptide levels during periods of intense reproductive competition, suggesting that the energetic cost of dominance increase with social challenges (Higham et al., 2011). Dominance can also involve changes in basal metabolism: dominant rainbow trout exhibit increased AMPK activity in skeletal muscle, supporting enhanced glucose uptake and fatty acid oxidation necessary for energetically demanding behaviors like patrolling and aggression (Gilmour et al., 2017). Likewise, dominant female rhesus macaques exhibit elevated levels of triiodothyronine (T3), consistent with increased metabolic demands (Jarrell et al., 2008). Subordinates also exhibit dynamic behavioral and physiological responses to their social position (Milewski et al., 2022). To avoid conflict, subordinates across species often closely monitor dominants and strategically inhibit their own aggressive behavior (Curley, 2016; Deaner et al., 2005; Desjardins et al., 2012; Pannozzo et al., 2007). and modify feeding patterns to minimize aggression, either avoiding high-ranking individuals altogether or, in some species, selectively choosing to feed alongside them to reduce aggression (Dale et al., 2017; Naud et al., 2016).

Previously we have shown that adult male outbred CD-1 mice reliably form linear social hierarchies (Lee et al., 2018, 2022; So et al., 2015; Williamson et al., 2016). Within these hierarchies, dominant males engage more frequently in patrolling, feeding, drinking, and scent-marking behaviors (Lee et al., 2018; Williamson et al., 2016). These animals also show increases in the production of major urinary proteins (MUPs), a metabolically costly class of molecules used in social status signaling (Lee et al., 2017a). To sustain their elevated output, dominant mice eat and drink more frequently (Lee et al., 2018) and exhibit metabolic changes including increased lipid catabolism, elevated energy production, and overall higher metabolic rates (Lee et al., 2022). Conversely, subordinate mice adjust their eating and drinking schedules to avoid dominant individuals, particularly those following aggression (Lee et al., 2018). These divergent behavioral and physiological phenotypes are companied by distinct patterns of neural gene expression (Lee et al., 2017b; Milewski et al., 2025; Milewski et al., 2025; Williamson et al., 2019; Williamson et al., 2017; Williamson et al., 2017).

Much of our previous research relied on live scoring of mouse behavior, a method that, while detailed, is inherently limited. Live observations are time-consuming, labor-intensive, and constrained by human availability and perceptual resolution. These limitations make it difficult to study the long-term expression of dominance behaviors, the fine-scale temporal dynamics of social hierarchies, or to differentiate behavior across light and dark cycles. To address these gaps, researchers have turned to automated systems such as video-based pose estimation and tracking software (e.g., EthoVision, DeepLabCut, SLEAP). However, these systems often require extensive training, consistent lighting, high-resolution video, and costly licenses (Geuther et al., 2019; Hong et al., 2015; A. Mathis et al., 2018; M. W. Mathis & Mathis, 2020; Noldus et al., 2001; Pereira et al., 2021) and still face difficulties when tracking animals in groups or in complex, three-dimensional environments (Panadeiro et al., 2021). An increasingly popular alternative to video-based tracking is the use of radio-frequency identification (RFID) systems, which have been widely used to monitor animal movements in natural and semi-natural settings including in mice (Dexter et al., 2016; Gelardi et al., 2019; König et al., 2015; Pereira et al., 2021; Rafiq et al., 2021; Scott et al., 2016; Vogt et al., 2024). RFID allows for continuous, non-invasive identification and location tracking of individuals across large time scales and under variable environmental conditions. Inferences can then be drawn about the spatiotemporal patterns of social interactions in groups of mice.

In the current study, we characterized how dominance hierarchies emerge and are maintained in complex social environments, with a particular focus on the energetic and daily activity consequences of social rank. We achieved this through the development and validation of an RFID-based system that continuously tracks the real-time behavior of group-housed male mice over extended periods. This automated approach enables fine-scale inference of dominance relationships based on movement and pairwise interactions, overcoming key limitations of traditional live-scored methods. By integrating this high-resolution tracking with hormonal analyses, we examine how dominant and subordinate roles shape access to resources, behavioral rhythms, and physiological states. Specifically, we test whether dominant individuals exhibit distinct patterns of activity across the light and dark phases and hormonal profiles consistent with increased energetic demands, and whether subordinates adaptively regulate their behavior in response to the social environment. This work not only provides a scalable, unbiased method for tracking social dynamics but also reveals how social status intersects with metabolic regulation and daily rhythms, offering a more comprehensive view of the physiological cost of dominance.

## Methods

### Subjects, Housing, and Experimental Design

7-8 week old male outbred CD-1 mice (N = 54) were obtained from Charles River Laboratory (Houston, TX, USA). Mice were initially housed in pairs in standard pine shaved bedding cages and were characteristically marked with a nontoxic animal marker (Stoelting Co., Wood Dale, IL, USA) for the purpose of consistent observer identification during social behavior scoring.

Pair social behaviors were scored during dyadic housing, and after three days, subjects were implanted with RFID tags (FBI Science; Viersen, Germany) assigned randomly into a multi-cage housing system comprised of 6 mice who were previously unfamiliar to each other (N = 9 groups). The transition to the multi-cage system occurred immediately before the shift from white light to red light. Each multicage system consisted of 6 standard mouse cages (7½” W x 11½” L x 5” H) inter-connected with Perspex tubes (diameter = 38mm), with food and water available *ad libitum* in one designated cage (Figure1A). All procedures were conducted with the approval of the Institutional Animal Care and Use Committee of the University of Texas at Austin.

### RFID Chip Implantation

Three days after arrival and pair housing, mice were implanted with RFID tags 1-2 hours before the onset of the dark cycle. Under isoflurane anesthesia, sterile 5mm biocompatible glass microchips (SO FDX-b-transponders; FBI Science; Viersen, Germany) were implanted subcutaneously with microchips in both their upper right thigh and dorsal neck between the shoulder blades. One day prior to group housing, mice were placed in the multi cage systems for 15 minutes to confirm that tags were detectable by all antennae.

### Behavioral Observations

In both pair housing and group housing agnostic behaviors (fighting, chasing, mounting) and subordinate behaviors (freezing, fleeing, subordinate posturing) were coded by trained observers and uploaded to a timestamped Google Drive survey via online link accessed via smartphone. Observers live scored social behaviors for 2 hours a day in the dark cycle (the active period for mice) for 10 days. In addition to live behavior scoring, RFID tags in mice were continuously read by circular antennas encircling the end of the tubes between all cages in the multi-cage system. These tag reads recorded the date and time (to millisecond accuracy) of every entrance and exit to all cages made by all individuals within each social group. The movement data output by the RFID was collected continuously throughout the 10 days of group housing using the IDE program (FBI Science, Germany). Utilizing both the live behavioral scoring and RFID data, linear social hierarchies were characterized. Hierarchical structure and stability were verified by calculating Landau’s modified h′ (de Vries, 1995) and David’s Scores (DS) of all individuals within the social network. DS is a measure of dominance based on the proportion of wins to losses adjusted for opponents’ relative dominance (Gammell et al., 2003). Individuals with positive DS are considered dominant and individuals with negative DS are considered subordinate.

### Blood Collection

Five days after arrival and two days after RFID tag implantation, blood was collected from all mice via submandibular blood draw. Blood was collected one hour after the onset of the dark cycle. Food and water were removed from all pair-housed mice approximately four hours before collection. On the tenth day of group housing, mice were sacrificed one hour into the dark cycle via decapitation following a four hour fast, and trunk blood was collected. For all blood samples, Sigma’s protease inhibitor cocktail (1mg/mL) and Roche’s Pefabloc SC (10mg/mL) were added to prevent protein degradation. All samples were collected in EDTA-coated Vacutainer tubes (Becton Dickinson) mixed by inversion, placed on ice for 45-60 minutes, then centrifuged to collect plasma. Samples were stored in −80°C fridge until processing.

### Hormone Multiplexing

A metabolic and pituitary hormone panel was established using samples taken from all mice that maintained subordinate or dominant status in pair and group housing conditions. Blood samples were collected by submandibular puncture before multi-chamber housing and by trunk blood collection on the last day of social group multi-chamber housing. For the hormone panel, samples from the highest- and lowest-ranked animals within each cohort were analyzed, resulting in 18 animals total (9 dominant and 9 subordinate). Plasma samples were processed with FlexMap Luminex bead arrays multiplexing at the Bioanalytic and Single-Cell Core (BASiC) at UT San Antonio Health. The commercially available multiplex panel was selected to simultaneously quantify metabolic and pituitary-associated hormones and inflammatory markers, including, Amylin (active), C-peptide2, Ghrelin (Active), MCP-1/CCL2, IL-6, Insulin, Leptin, PPY, Secretin, and TNF-alpha using a Millplex kit (MMHMAG −44K, MilliporeSigma). Data that did not meet predefined assay quality-control criteria were excluded from analysis.

### Statistical Analysis

All analyses and visualization were performed using R Statistical Software (v4.3.2; R Core Team 2023). For both the observational and RFID tracking data we used the ‘competè R package (Curley, 2016) to create sociomatrices of dominance behaviors, calculate directional consistency, non-normalized David’s Scores (Gammell et al., 2003). Glicko Rankings were calculated to measure temporal dominance rankings using the ‘playerRatings’ R package (Stephenson and Sonas., 2012; Glickman, 1999).

To determine dominance events using RFID data we identified when one mouse chased/followed another mouse rapidly through tubes in the cage system. A tube transition is defined as a movement from one end of a tube (flagging the first antenna traveled through as the entrance point of the mouse) to the other (flagged as the second antenna traveled through as the point of exit). Using the entrance and exit time recordings, we determined the directional movement of a mouse and classified a follow as a point when another mouse performed the same tube transition within a recent time window. We identified a follow as a “chase” by using a 250ms window because interactions flagged within that window demonstrated the highest directional consistency of chases (**Supplemental Figure S2**).

The overall activity of each mouse was classified as the number of tube transitions performed, the length of time between tube transitions on average, and the number of chases performed. We defined quiescence as the length of time in between tube transitions for each mouse. All statistical comparisons were performed using generalized linear mixed effect models (GLMMs) in order to control for cohort as a random effect. These linear models were built using the lmer and glmer functions from the lme4 library (Bates et al., 2023). To visualize the distribution of outcome variables (to check for normality) the moments library was used to report skewness values of outcome variables (Komsta, 2005). If the coefficient of skewness exceeded 0.3 for any outcome variable, more right skewed distributions were applied in glmer (using the Gamma distribution). If the coefficient of skewness was less than −0.3 for any outcome variable, outcome variables were squared and a gamma distribution was applied if the squared transformation did not sufficiently normalize the outcome variables. If the outcome variable distribution was between −0.3 and 0.3, the lmer function was used. To calculate the association between non-normalized David’s Score (DS) from observer coded agonistic interactions and non-normalized David’s Score (DS) from automated follow tracking we built a linear mixed effect model with agonistic DS predicting automated DS with cohort as a random effect.

To evaluate spatial dynamics of different ranks in the housing system, we grouped RFID antennas into “zones” surrounding each cage (see **Figure 1A**). We then calculated the ratio of transitions each mouse made in each zone by dividing the number of transitions made within that zone by the total number of transitions they made. We then built a mixed effects model predicting the outcome of proportion in each zone with the fixed effect of non-normalized DS and random effect of cohort. Another spatial use measure we included was the amount of time spent in each zone, determined by defining whether the most recent transition placed a mouse within a particular zone and recording that as time they were in that zone until they make another transition that puts them in a new zone. Then we built a mixed effects model predicting the outcome of time spent in each zone with the fixed effect of non-normalized DS and random effect of cohort.

**Figure 1:**
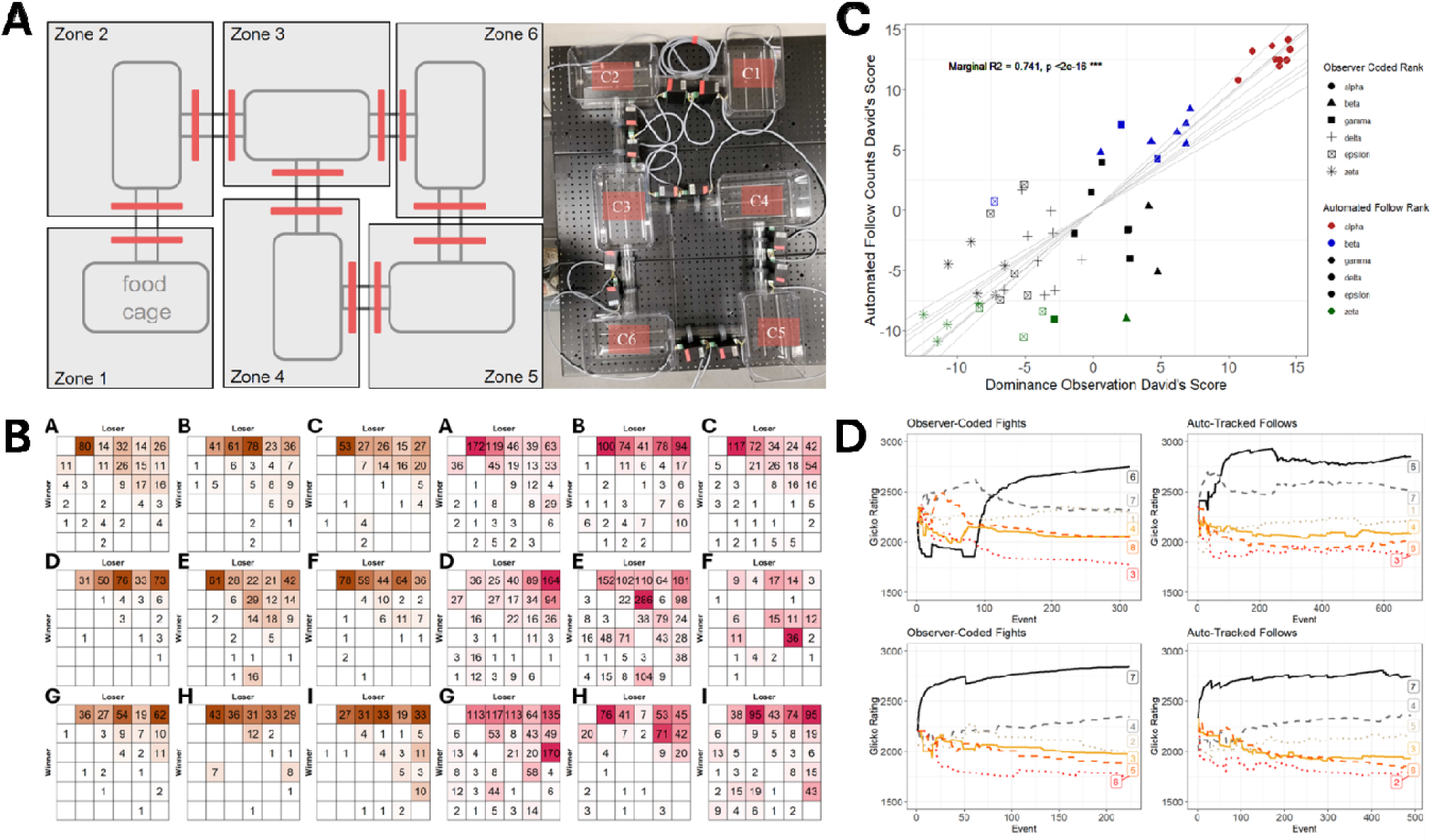
A) The multi-cage RFID housing system, separated into zones. Mice are first introduced to the RFID housing system by being placed in the food cage (C1, zone 1) right before the transition from white light (inactive phase) to red light (active phase). Thick red lines on the plot represent the antennas that the mice must move through to transition through the tubes to get from cage to cage. **B) Left: Win-loss sociomatrices based on observer coded aggression data.** Results are presented for all 6 mice in cohorts A-I. **Right: Win-loss sociomatrices based on automated follow data.** 250ms window dyadic follow results are presented for all 6 mice in cohorts A-I. **C): Relationship between David’s Scores in agonistic observations vs automated follows.** Alpha males were consistently identified in all nine cohorts between agonistic interactions and automated follow hierarchies. Two out of seven beta males were also consistently identified. The relationship between David’s Score (opponent-adjusted win proportion of each individual) in agonistic interactions and automated follows is significant such that a higher David’s Score in dominance observations (indicating greater dominance) corresponds to a higher David’s Score in automated follow interactions. The grey lines in the plot represent the lines of best fit for each of the nine cohorts. **D) Dominance (Glicko) ratings of male mice within cohorts A (top panel) and G (bottom panel) across behavioral events (automated follows vs. agonistic interactions).** Automated follow tracking identified the cohort A alpha as high-ranking early and as the top-ranked male within 100 dyadic events, whereas observer-scored agonistic interactions initially classified him as low-ranking. In cohort G, both methods immediately identified the alpha male, while gamma and delta ranks remained contested. The evolution of dominance ratings across time for all cohorts is shown in **Supplemental** Figure 4.

Using the measure of time spent in a zone, we calculated overlaps of time spent in each zone between mice to create an association proportion. This proportion estimates the amount of time a mouse spends with a mouse of a different rank in their cohort of all 10 days of recording. To compare association proportions between each rank, we calculated the average association proportion for each rank and created a linear model predicting the association proportion by rank with the random effect of cohort. Then, we collected the p values for each of the comparisons in the model, i.e. the focal rank with each of the other ranks, for all nine cohorts and randomly shuffled the rownames/colnames of each of the nine matrices into new matrices. Then we recalculate the mean association indices for each rank and then collect the p-values from a new linear model predicting the association proportion by rank with the random effect of cohort in these shuffled averages. We repeat this process for 1000 permutations, and then calculate the number of times out of 1000 where the p value for each of the rank pair comparisons were lower or equal to the observed we originally calculated. This was repeated with each rank (alpha-zeta) as the focal rank.

To build the hormone panel analysis models, we used the lmer and glmer function to predict post-group housing hormone concentrations. The use of lmer or glmer was determined by analyzing the normality of the outcome variable of post group housing hormone concentration. This was calculated for each hormone in the panel. The only hormone that was certainly non-normal was Secretin, which had a floor effect in 8 out of 17 subjects, and analysis was better suited to a Poisson distribution than a normal distribution. The Poisson distribution was applied by using glmer and setting the family to poisson. All predictor and outcome variables were rounded to whole values to suit the count data expectation when utilizing a Poisson distribution. For the other normally distributed outcome variables, lmer models were built. Fixed effects within the model included the pre-group housing ranks of each subject as well as their pre-group housing hormone concentrations, and proportion of activity made in the active phase, calculated by computing the number of transitions made in the active phase (red light) divided by the total number of transitions made in both the inactive phase (white light) and active phase (red light).

When including rank in models, we used pre-group housed (pair-housed) ranks as a binarized predictor. Rank prior to group housing was included over post-group housing rank because it contributed more value to models; improving fit and interacting with other variables more substantially than post-group housing rank. We included pre-group housing hormone concentrations to control for individual base hormone concentration as well as to explore if individuals showed consistency in hormone levels before and after group housing. Finally, we calculated the proportion of each mouse’s total transitions that occurred during the red-light active phase to use as a predictor. Because this measure was not significantly correlated with pre-group housing rank, unlike total activity, it could be included alongside rank in the hormone models to assess whether the temporal distribution of activity across the light-dark cycle was independently associated with post-group housing hormone concentrations.

## Results

### Automated RFID tracking recapitulates highly linear dominance hierarchies

All 9 groups of six males formed highly linear dominance hierarchies. **Figure 1B** shows the win-loss sociomatrices based on behavioral observations of all agonistic interactions. Each group had a significant directional consistency (DC) (median[iqr]: 0.95[0.89,0.98], all p<0.001), indicating that dominance behaviors were largely directed from more dominant males to more subordinate males. Within each group it was possible to assign individual ranks to males based on David’s Scores (DS) (**Supplementary Figure 1**). All alpha (rank 1) males had the highest DS above 0 indicating that they win more contests than they lose against all other individuals. Across groups, all beta (rank 2) males and 6 out of 9 gamma (rank 3) males exhibited positive DS, indicating that these males win more contests on average than they lose and are therefore sub-dominant. Subordinate males had DS below 0.

Win-loss sociomatrices were also constructed based on chases through tubes determined from RFID tracking data (**Figure 1B**). We considered a chase to be when one mouse followed another mouse through a tube within 250ms of the first mouse passing through the tube. Groups ranged between 151 and 1570 total number of chases per group over the 10 days of social housing (median[iqr]: 555[404,706]). All groups had significant linear hierarchies with a significant DC (median[iqr]: 0.83[0.78, 0.87], all p<0.001), with DC being significantly higher in agonistic sociomatrices compared to RFID chasing sociomatrices (**Supplemental Figure 1C**, β=-0.11 ± 0.03, p<0.01). The DC of RFID tracking sociomatrices was maximal with window sizes of 250ms and decreased with increasing window sizes (**Supplemental Figure 2**), indicating that shorter windows are most sensitive to identifying chases. However, it should be noted that even using window sizes of 1s it is possible to classify the individual ranks and hierarchical structure of each group.

The high linearity of hierarchies is also evidenced by the remarkably few inconsistencies in the direction of relationships. Inconsistent relationships are defined as where a pair of individuals in a linear dominance hierarchy contradict their rank order with a lower ranked member winning more competitions between themselves and a higher ranked group member than they lose.(**Supplemental Figure 3A**). Out of 15 total relationships between 6 mice in each cohort, there was an average of 0.22 inconsistencies in relationships in agonistic sociomatrices compared to an average of 1.89 inconsistencies in RFID tracking sociomatrices using data aggregated across the whole 10 days (**Supplemental Figure 3B**).

We found that the rank orders of individuals determined by behavioral observations and RFID tracking were remarkably consistent (**Figure 1C**). Dominant alpha males are reliably identified with both measures with the same alpha male being identified in all nine groups. We found that the relationship between DS based on behavioral observations and RFID tracking was highly significant (F(1,44) = 151.58, p < 0.0001). It is possible that the greater number of inconsistencies in RFID tracking sociomatrices and less consistent classification of lower ranks between observer coded fighting and automated follow tracking is due to the greater volume of data collected in RFID detection (mean ± SE: 685.0 ± 140.0 chases vs. 269.2 ± 16.9 agonistic interactions) and the additional observations that are collected during white light in RFID detection. Indeed, when comparing the development of hierarchy formation (**Figure 1D & Supplemental Figure 4**) automated tracking more rapidly identifies the alpha and beta males as they rise than observer coded follows, although both methods identify the stabilization of ranks over time (**Supplemental Figure 5**).

### Alphas exhibit greater activity and reduced quiescence than lower-ranking males

We used automated tracking data to determine how the activity levels of mice varied by social rank. Alphas males show significantly greater activity (in the form of tube transitions made) than any other rank of male within the hierarchy (comparison: β ± SE; alpha-beta: β=-0.96 ± 0.21, p < 0.001; alpha-gamma: β=-0.65 ± 0.21, p = 0.002; alpha-delta: β=-0.76 ± 0.22, p < 0.001; alpha-epsilon: β=-0.77 ± 0.21, p < 0.001; alpha-zeta: β=-0.74 ± 0.21, p < 0.001). These significant differences emerge within the first hour of group housing on day 1(**Figure 2A**). Similarly, alphas show significantly lower maximum quiescence durations (time spent inactive) than their lower ranking counterparts (**Figure 2B**, comparison: β ± SE; alpha-beta: β=0.39 ± 0.04, p < 0.001; alpha-gamma: β=0.42 ± 0.04, p < 0.001; alpha-delta: β=0.40 ± 0.04, p < 0.001; alpha-epsilon: β=0.43 ± 0.04, p < 0.001; alpha-zeta: β=0.46 ± 0.04, p < 0.001). Indeed, the average time inactive between tube transitions for alpha males in red light was only 18 seconds (median[IQR]: 1.3s[0.0,8.3]) and 34 seconds in white light (median[IQR]: 1.5s[0.0,8.4]). All other ranks average 36 seconds inactive in red light (betas: 1.0s[0.0,8.9]; gammas: 1.1s[0.0,10.2], deltas: 1.1s[0.0,9.8], epsilons: 1.3s[0.0,11.6], zetas: 1.4s[0.0,11.8]) and 66 seconds inactive in white light (betas: 1.3s[0.0,8.8]; gammas: 1.5s[0.0,10.6], deltas: 1.6s[0.0,11.0], epsilons: 1.6s[0.0,11.5], zetas: 1.6s[0.0,9.2]). Day also significantly predicted the maximum inactive period each day (quiescence), with greater maximum quiescence increasing over days (χ2(1,10) = 13.2, p < 0.001). As expected, significantly greater maximum quiescence values were observed for all mice during the white light (inactive) phase than the red light (active) phase (χ2(1,10) = 785.5, p < 0.001).

**Figure 2:**
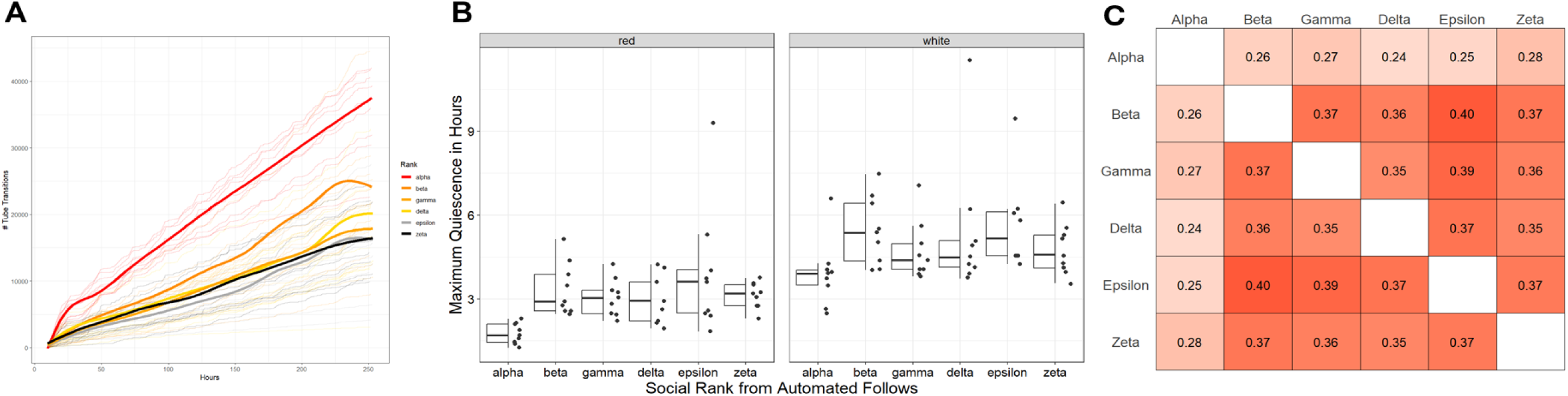
A) Activity (number of tube transitions) for all males, with average number of tube transitions per hour colored by rank. Alpha males show consistently greater numbers of tube transitions than other males within their hierarchy. **(B) Maximum quiescence (time spent inactive between transitions) between ranks in across light phases.** Alpha mice show significantly lower quiescence (indicating higher activity levels) compared to all other ranks. Lower ranked mice show no significant differences in quiescence compared to each other. **C) Average association index across all cohorts by rank.** Alpha males show significantly lower average association proportion to other mice than Beta-Zeta males show with each other. Deeper red in the figure indicates higher association proportion.

### Higher-ranked males spend less time with lower-ranked individuals, who occupy more peripheral zones

To investigate spatial usage, we separated the RFID setup into six zones corresponding to the six cages and the surrounding tubes/antennae of those cages (**Figure 1A**). We found that higher ranking males show a significantly greater proportion of transitions in zone 3 (β=0.008±0.004, p = 0.022) and approach a significantly greater proportion of transitions in zone 1 (β= 0.0007 ± 0.0004, p = 0.069) compared to lower ranking males (**Supplemental Figure 6**). In contrast, lower ranking males show greater proportion of transitions in zone 5 (β=-0.007±0.003, p = 0.009) and approach a significantly greater proportion of transitions in zone 6 (β=-0.004 ± 0.002, p = 0.096) compared to the proportion of transitions shown by higher ranking mice (**Supplemental Figure 6**). Time spent in each zone is also significantly related to automated rank, with higher ranking males showing significantly more time spent in zone 3 (β=1.584e+06 ± 5.616e+05, p = 0.005) and zone 4 (β=0.020 ± 0.009, p = 0.022) and lower ranking males showing significantly more time spent in zone 6 (β=-0.012 ± 0.005, p = 0.027) (**Supplemental Figure 6**). Compared to other mice, alpha males associate significantly less (based on time spent in the same cage) with mice of other ranks (all alpha associations p<0.005 after permutation comparison) (**Figure 2C**).

### Housing, rank and daily activity and are associated with circulating plasma hormone concentrations

Using hormone multiplexing, plasma hormone concentrations were analyzed at two timepoints: i) before social group housing during pair-housing and ii) on the final day of social group housing. We determined the plasma concentrations of C-Peptide, Insulin, Leptin, Ghrelin, Interleukin-6, Monocyte Chemoattractant Protein-1, Secretin, Peptide YY, Tumor necrosis factor alpha, and Amylin at each timepoint. To evaluate the influence of social rank and activity on these hormones, we built a model that incorporated i) binarized rank in pair housing prior to group housing; ii) hormone concentration prior to group housing; iii) proportion of transitions made during the red-light phase versus the white-light phase as a measure of activity. For all hormones investigated, except for C-peptide and Insulin, hormone concentrations were positively associated across the two time points (**Figure 3A**). Dominance rank in pre-group housing (pair housing) was associated with lower plasma C-Peptide, Insulin, and Secretin (**Figure 3B, Supplemental Figure 7 and 8**). Higher proportion of activity in the red-light period was also associated with lower post-group housing plasma Secretin (**Figure 3C**). Pre-group housing dominance status led to a greater positive relationship between proportion of activity in red light and post-group housing hormone concentration in interleukin-6, monocyte chemoattractant protein-r1, and peptide Y (**Figure 3D**). For all significance, model estimates, and standard errors see **Supplemental Figure 7**.

**Figure 3:**
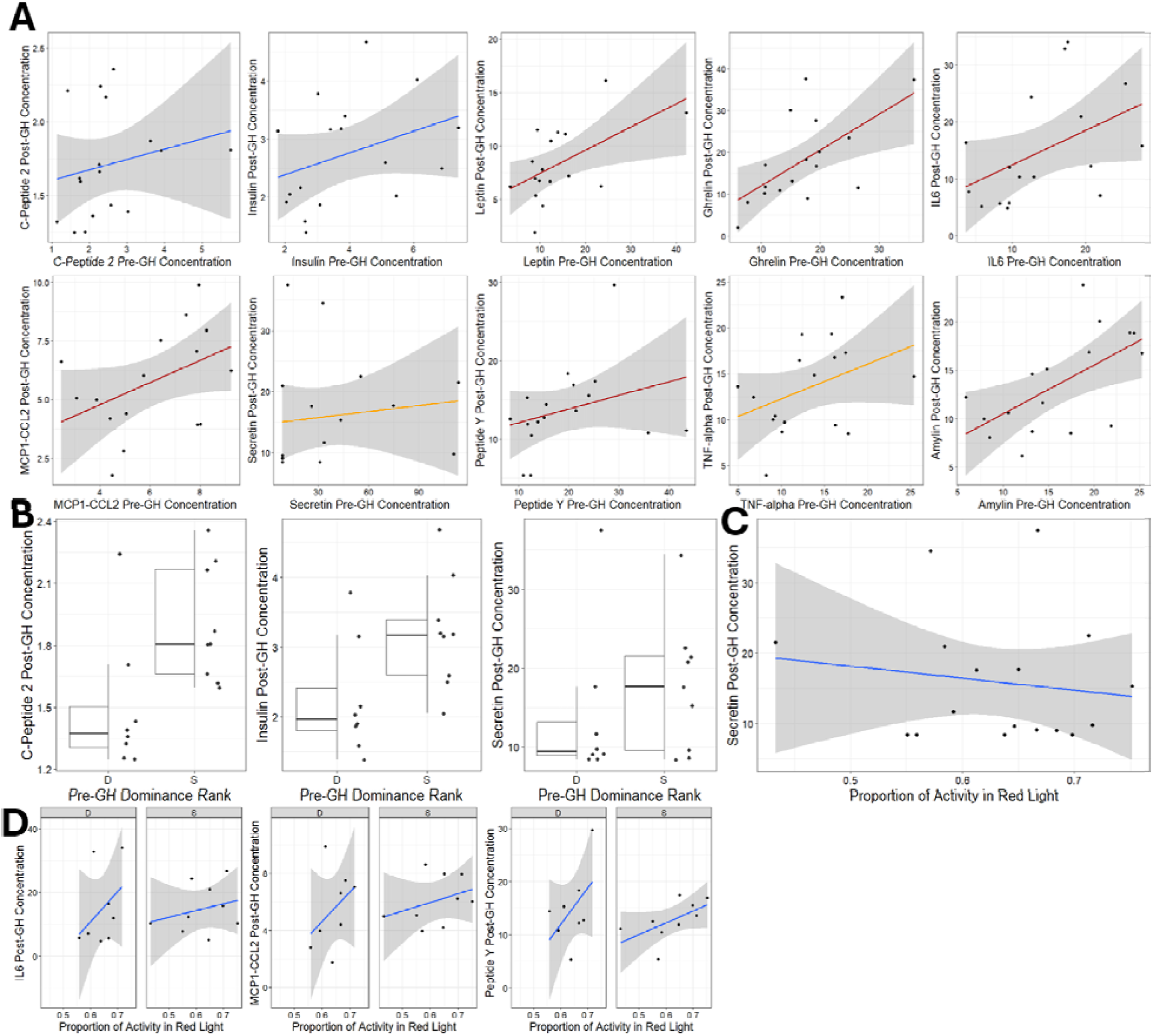
A) Pre-group housing hormone levels significantly predict post-group housing hormone levels in most metabolic hormones measured. The relationship between the pre-group housing hormone levels (x axis) and the post-group housing hormone levels (y axis) is presented. Red colored lines represent significant predictive relationships (p < 0.05). Orange colored lines represent marginally significant predictive relationships (p < 0.1). Only C-peptide and Insulin pre-group housing hormone levels did not significantly predict post-group housing hormone levels. All significant relationships were positive, such that higher pre-group housing hormone levels were predictive of higher post-group housing hormone levels. **B) Pre-group housing rank significantly predicts post-group housing concentrations of C-Peptide, Insulin, and Secretin.** Mice that were dominant in their pre-group housing pairs demonstrate significantly lower concentrations of these three hormones after ten days of group-housing. **C) Proportion of activity in red-light versus all activity during group-housing significantly predicts Secretin concentration after group-housing.** This relationship is such that mice that showed a relatively higher proportion of transitions during red light have lower concentrations of secretin at the end of the group-housing period. **D) Pre-group housing rank interacts with active-phase activity to predict IL-6, MCP-1, and PYY concentrations.** For all three hormones, the relationship between the proportion of activity occurring during the red-light phase and post-group housing hormone concentration was stronger in pre-group housing dominant mice.

## Discussion

In this study, we used continuous RFID tracking and hormonal analyses to investigate the emergence, maintenance, and physiological correlates of dominance hierarchies in group-housed male mice. Our results demonstrate that male CD-1 mice rapidly form highly linear social hierarchies, with dominant individuals exhibiting elevated activity levels, reduced quiescence, and distinct patterns of spatial use. These behavioral differences were mirrored by rank-related differences in metabolic hormone concentrations, with dominant males showing hormonal profiles consistent with greater energetic demands. Subordinates, in contrast, displayed lower activity levels and greater use of peripheral areas, patterns that may be consistent with conflict avoidance and energy conservation. Automated RFID tracking provided a scalable and temporally precise alternative to live behavior scoring, allowing us to detect early rank emergence and assess social dynamics across the full light-dark cycle. Together, these findings highlight the behavioral and energetic consequences of dominance, as well as rank-related differences in dark-light activity and establish RFID-based tracking as a powerful tool for studying the interplay between social structure, behavior, and physiology.

By developing a real-time RFID tracking system for use in a multi-cage social housing environment, we were able to continuously monitor the movements and interactions of group-housed CD-1 male mice over a 10-day period. This system tracked behavior across both the light (inactive) and dark (active) phases and inferred dominance-related interactions, specifically chases, from short-latency tube-following events. These automated data were compared directly with manually scored social behaviors, including aggression and submission, enabling us to validate dominance hierarchies based on follow behavior. Previous RFID approaches have been used successfully to quantify migration, crossings, general movement, proximity-based social networks, and resource use in wild and semi-natural settings (Barlaup et al., 2018a, 2018b; Gelardi et al., 2019; König et al., 2015; Rafiq et al., 2021; Renardy et al., 2020; Ross et al., 2022; Scott et al., 2016; Testud et al., 2019). However, many such systems rely on more widely spaced sensors or co-location measures and therefore provide relatively coarse measures of movement or social association (Dexter et al., 2016), making them less suited to distinguishing rapid dominance-related interactions. More recent lab-based systems, such as ColonyTrack (Doludda et al., 2023) and Eco-HAB (Chen et al., 2025), have used tightly spaced antennae to estimate “chases” from sequential passes through a shared tube within short time intervals. Our system builds on this approach by incorporating two antennas per tube, allowing us to infer movement direction and exclude ambiguous follows. Unlike studies that rely on tube tests alone for dominance validation, we used full-system live coding of agonistic and subordinate behaviors, including fighting, fleeing, and subordinate posturing, providing a broader ethological assessment of rank (Lindzey et al., 1961) and allowing direct comparison with RFID-derived rankings. We also used a stricter 250ms follow threshold than the 1s windows used in ColonyTrack and Eco-HAB. Shorter follow windows produced more directionally consistent hierarchies, supporting the use of the 250ms threshold for identifying dominance-related follows. Using this method, we observed highly linear dominance hierarchies across all nine cohorts, with particularly strong agreement between automated and observer-based methods in identifying alpha males. As in prior studies, lower-ranking positions were somewhat more variable, particularly between methods (Williamson et al., 2016, 2019). This variability may reflect the greater volume and temporal coverage of RFID data or may indicate that RFID captures additional status-related behaviors, such as persistent spatial displacement or patrolling, that may be missed during shorter observation windows.

Dominant males exhibited behavioral and endocrine profiles consistent with greater energetic demands associated with maintaining high social status. We found that alpha males exhibit significantly greater overall activity, as measured by tube transitions, reduced periods of quiescence, and distinct patterns of spatial use. They also spent less time in close spatial association with lower-ranking males. These results are consistent with previous findings that dominant mice engage in more extensive patrolling and territorial behavior, increased feeding and drinking and reduced sleep (Audet et al., 2011; Bartolomucci et al., 2001; Karamihalev et al., 2019; Lee et al., 2017b, 2018; Williamson et al., 2016). These behavioral patterns were paralleled by distinct endocrine profiles. Specifically, after accounting for pre-group housing hormone concentrations and the proportion of activity occurring during the red-light active phase, higher pre-group housing dominance rank predicted lower concentrations of insulin and C-peptide. Insulin and C-peptide provide related indices of metabolic state, with C-peptide serving as a marker of endogenous insulin secretion (Luzi et al., 2007). The lower concentrations of C-peptide and insulin in dominant males are consistent with previous findings that high-ranking individuals in highly competitive social contexts can experience greater energetic demands and negative energy balance. For example, high-ranking male chimpanzees and rhesus macaques show reduced C-peptide levels during periods of heightened reproductive competition or social challenge associated with increased energetic expenditure (Thompson et al., 2009; Higham et al., 2011).

In contrast to dominant males, subordinate individuals displayed behavioral profiles consistent with reduced energetic expenditure and possible conflict avoidance. Lower-ranked mice made fewer transitions between zones, spent more time in peripheral cages farther from the food source, and exhibited longer periods of quiescence. These patterns may reduce encounters with dominant animals and are consistent with previous work showing that subordinates can suppress overt aggression, alter feeding schedules, and closely monitor dominants to avoid conflict (Curley, 2016; Deaner et al., 2005; Naud et al., 2016). Subordinates also showed a greater difference in quiescence between the active and inactive phases, suggesting stronger light-dark differentiation in inactivity. Social stress can also alter energy balance, with chronic social defeat studies reporting reductions in food intake, feed efficiency, or body weight in susceptible animals, although these effects vary by strain and context (Iio et al., 2011; Krishnan et al., 2007; Okamura et al., 2022; Razzoli et al., 2011; Sial et al., 2016; Smith et al., 2023; Toyoda, 2017). Together, these findings suggest that subordinate status is associated with a lower-activity behavioral phenotype that may reflect reduced energetic expenditure and avoidance of costly social interactions.

A notable finding from our hormonal data is the high degree of individual consistency in hormone concentrations across the transition from pre-group to group housing. For nearly all hormones measured, excluding C-peptide and insulin, pre-group housing levels strongly predicted post-group housing levels, suggesting that individuals maintain relatively stable endocrine profiles despite substantial changes in their social environment. This pattern supports the idea that metabolic hormone regulation may be shaped by trait-like physiological baselines, rather than being entirely driven by social rank or context (Cohen & Hamrick, 2003, 2003; Holtmann et al., 2017). Indeed, one previous study in socially housed primates also found that circulating stress hormone levels often return to individual set-points after an initial adjustment to social conditions, highlighting the stability of internal physiological states across environmental transitions (Czoty et al., 2009). Insulin and C-peptide were notable exceptions to this general pattern, with pre-group concentrations not significantly predicting post-group concentrations. Their similar patterns are physiologically consistent with their coupled secretion, as C-peptide is released alongside insulin during proinsulin processing (Luzi et al., 2007). Notably, these were also the hormones most clearly associated with dominance rank, suggesting that insulin and C-peptide may be particularly responsive to changes in social and energetic context.

Beyond the rank-related effects on insulin and C-peptide, several other hormones were associated with the temporal distribution of activity across the light-dark cycle. Post-group housing secretin concentrations were lower in mice that were dominant prior to group housing and in individuals with a greater proportion of their total activity occurring during the red-light active phase. Given evidence that secretin signaling is involved in sleep and circadian regulation (Sugiyama et al., 2022), these findings suggest that secretin may be related to the temporal organization of activity in socially housed mice. For PYY, the association between pre- and post-group housing concentrations varied according to red-light activity. Because circadian disruption can alter satiety signaling, including PYY (Martchenko et al., 2020; Moghadam et al., 2017), these results suggest that the timing of activity may be relevant to metabolic hormone regulation. We also observed more limited effects for inflammatory markers. Previous work from our group found that socially subordinate animals show a higher proportion of peripheral innate immune cells, increased proinflammatory gene expression in liver and spleen, and upregulation of immune-related pathways in these tissues (Lee et al., 2022). In the present study, however, post-group housing IL-6 and MCP-1 concentrations were predicted only by interactions between pre-group housing dominance status and the proportion of activity occurring during the red-light phase, and these effects were small. This discrepancy may reflect differences in the timing of immune activation or indicate that circulating cytokine concentrations do not fully capture persistent tissue-level inflammatory changes. Overall, these findings suggest that, beyond the more stable endocrine profiles observed for many hormones, specific metabolic and immune markers are associated with both social status and the temporal organization of activity across the light-dark cycle.

In conclusion, we developed and validated an automated RFID-based system for tracking social dominance in group-housed mice, demonstrating high consistency with observer-coded hierarchies. This approach provides continuous, behavioral data across light and dark phases, enabling the detection of temporal dynamics and spatial patterns that traditional scoring methods often miss. Our findings reveal that dominant and subordinate males differ in activity, quiescence, spatial use and metabolic hormone profiles, and that many hormone concentrations remain remarkably stable across changes in social context. We further show that both social status and the temporal distribution of activity across the light-dark cycle are associated with specific hormonal outcomes, linking social organization with metabolic physiology. Moving forward, this system offers a scalable framework to test theory-driven questions about dominance hierarchy emergence, and how social organization evolves in larger or more complex groups.

## Data Availability Statement

All data and code associated with this study are publicly available at the following GitHub repository: https://github.com/jalapic/seese_rfid.

## Acknowledgements

We wish to thank Dr. Frank Buschmann from FBI Science, Viersen, Germany, for his assistance in setting up the RFID system.

## Notes

https://github.com/jalapic/seese_rfid

### Competing Interest Statement

The authors have declared no competing interest.

