## Supplemental Figures for "Socially dominant male mice in social hierarchies identified via automated RFID tracking exhibit elevated activity levels and circulating markers of higher metabolic demand"

**Supplemental Figures and Tables**

**Supplemental Figure 1**

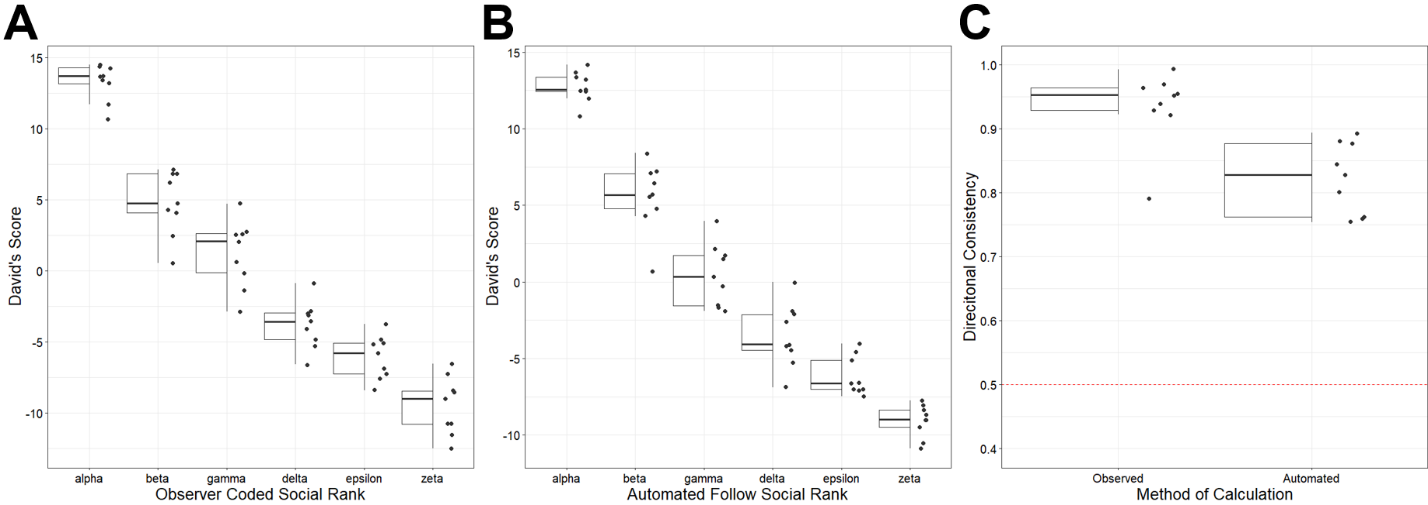

**Supplemental Figure 1: (A) Non-normalized David’s Scores for all social ranks in 6-mouse social groups.** David’s Scores as determined by observer coded agonistic interactions are presented for all mice across 9 cohorts. Alpha and Beta ranks are all above 0, and a majority of Gamma ranks are above 0 (6 out of 9 gammas). **(B) Non-normalized David’s Scores for all social ranks in 6-mouse social groups.** David’s Scores as determined by automated follow interactions are presented for all mice across 9 cohorts. Alpha and Beta ranks are all above 0, and a majority of Gamma ranks are above 0 (5 out of 9 gammas). **(C) Directional consistency values for all 9 cohorts when calculating hierarchy with agnostic interactions (Observed) or automated follow data (Automated).** Observer coded agonistic interaction directional consistency values for all nine mouse cohorts are compared to 250ms window automated dyadic follow directional consistency values. The horizontal red dotted line on the plot at 0.5 directional consistency represents complete unidirectionality of a sociomatrix, such that dominant and subordinate individuals are equally likely to win all contests (ie random chance).

**Supplemental Figure 2**

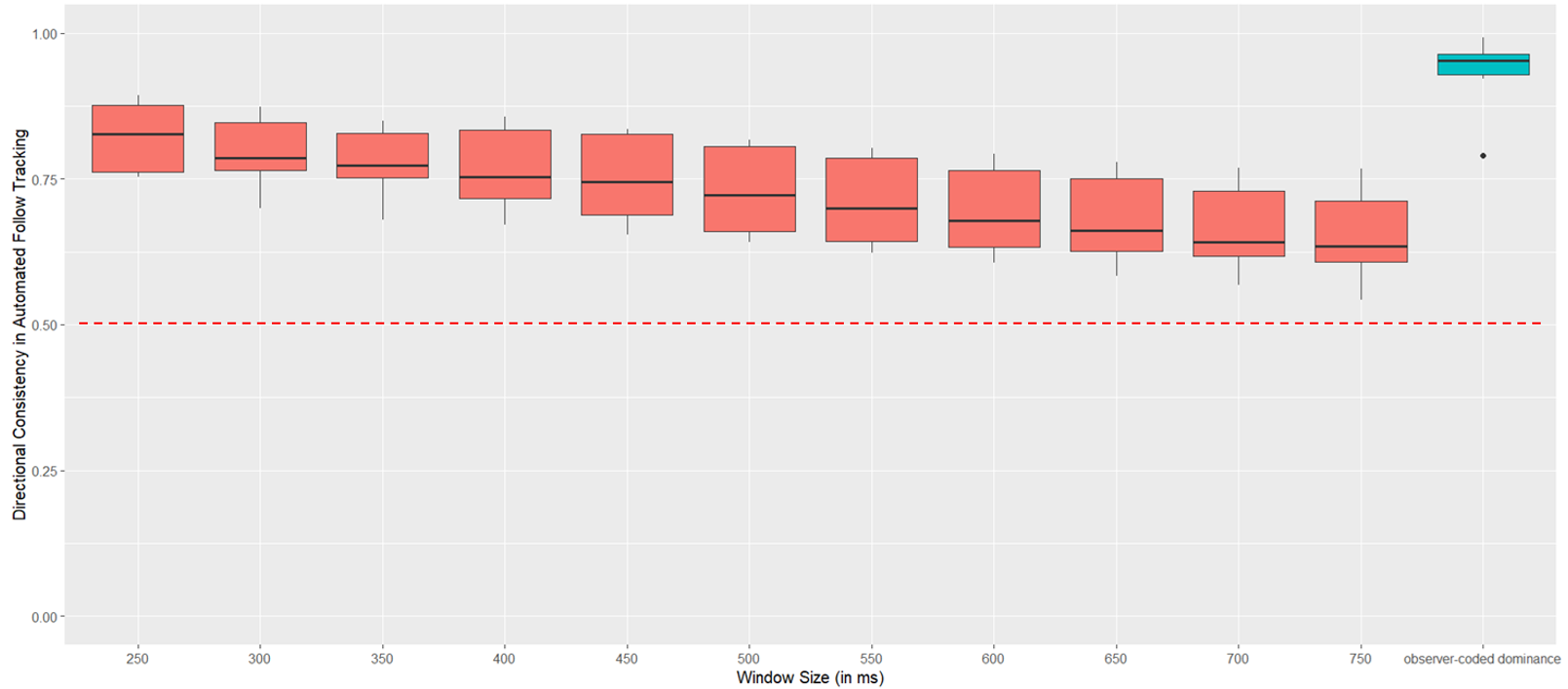

**Supplemental Figure 2: Directional consistency of metrics for different window sizes (red filled boxplots) compared to observer coded agonistic interaction directional consistency (blue filled boxplot).** Boxplots are presented for 11 different window sizes tested for the cut off of what can be considered a “chase” or likely aggressive interaction using the automated RFID system. Window sizes from 250 to 750 ms were used. As window sizes, decrease the number of “chases” identified decreases but the directional consistency of chases increases. Because the 250 ms window produced the highest directional consistency, most closely matching observer-coded dominance patterns while still capturing a sufficient number of chases, it was selected for automated dominance tracking. The red dotted line on the plot at a DC value of 0.5 represents complete unidirectionality of a sociomatrix, such that dominant and subordinate individuals are equally likely to win all contests (ie random chance). No window size fell below this minimum DC value.

**Supplemental Figure 3**

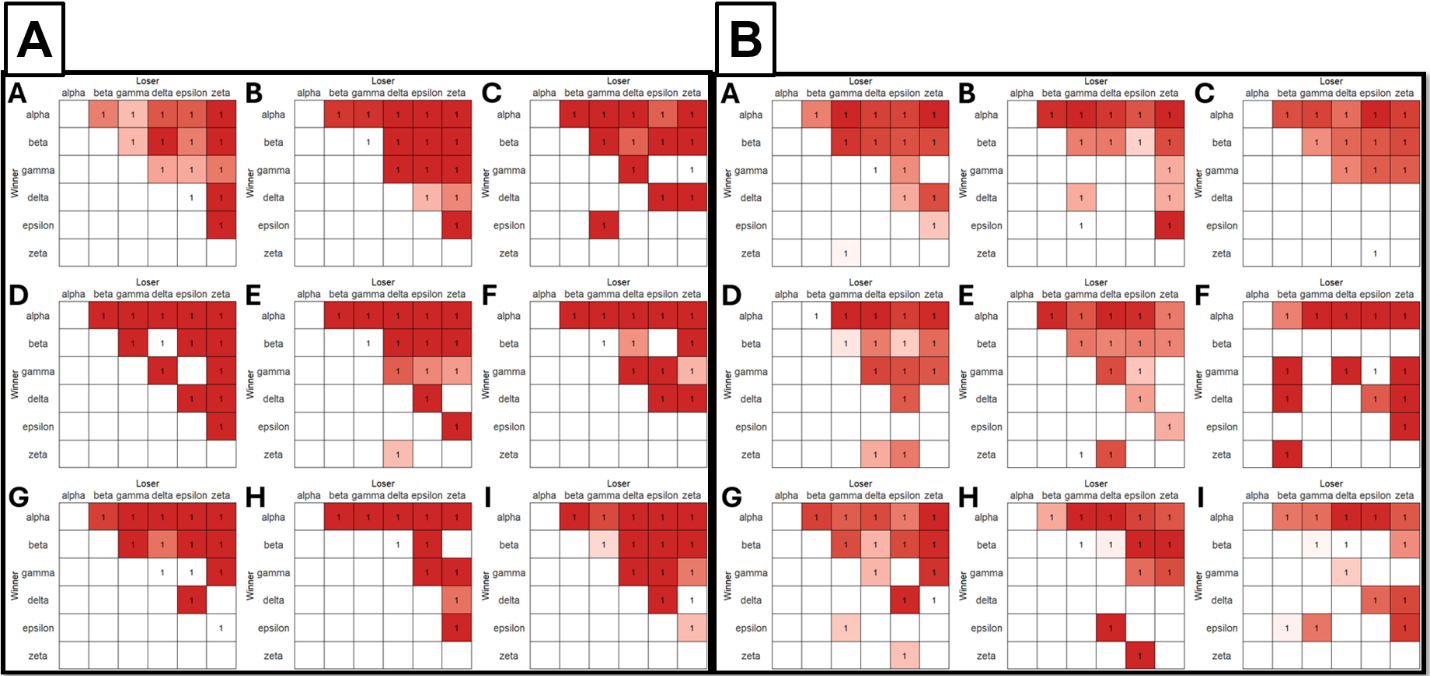

**Supplemental Figure 3: (A) Inconsistency matrices for observer coded agonistic interactions.** These matrices are organized based on rank, with highest ranking mice in the leftmost columns and top rows of each matrix. Inconsistencies appear when the individual in a lower row is a winner over the individual in the column. Only 1 inconsistency is observed in Cohort C and Cohort E in observer coded agnostic matrices. Cohort letter is labeled in the upper left corner of each sociomatrix. **(B) Inconsistency matrices for automated follows.** These matrices are organized based on rank as determined by automatically detected “chases”, with highest ranking mice in the leftmost columns and top rows of each matrix. Inconsistencies appear when the individual in a lower row is a winner over the individual in the column. Cohort F demonstrates 3 inconsistencies, cohorts B,D,E,G,H, and I show 2 inconsistencies, and cohorts A and C show 1 inconsistency each. Cohort letter is labeled in the upper left corner of each sociomatrix.

**Supplemental Figure 4**

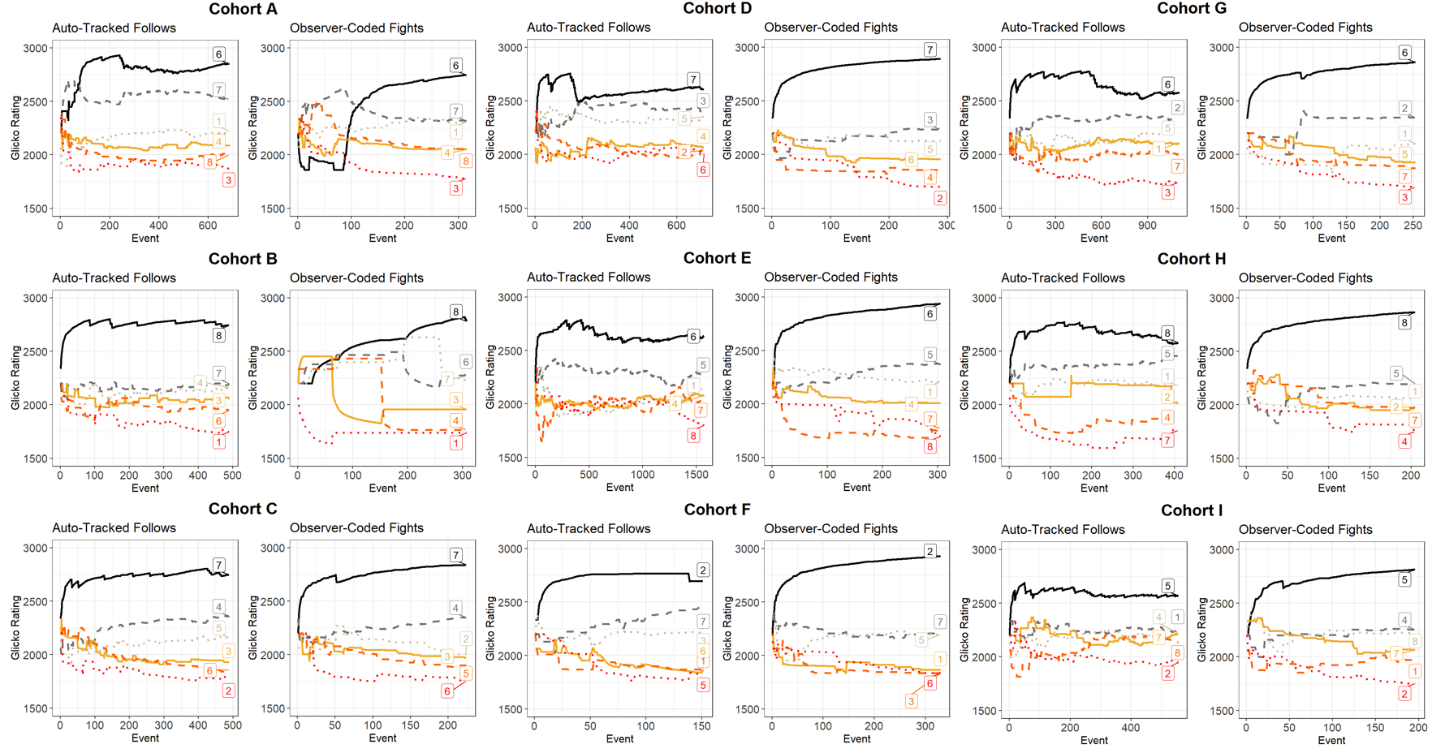

**Supplemental Figure 4: Evolution of Dominance (Glicko) ratings after each agonistic interaction for all cohorts presented for both auto-tracked rankings from RFID chases and agonistic observer coded interactions.** Rank progressions demonstrate the same alpha emerging in all cohorts across both auto-tracked and observer-coded hierarchy tracking.

**Supplemental Figure 5**

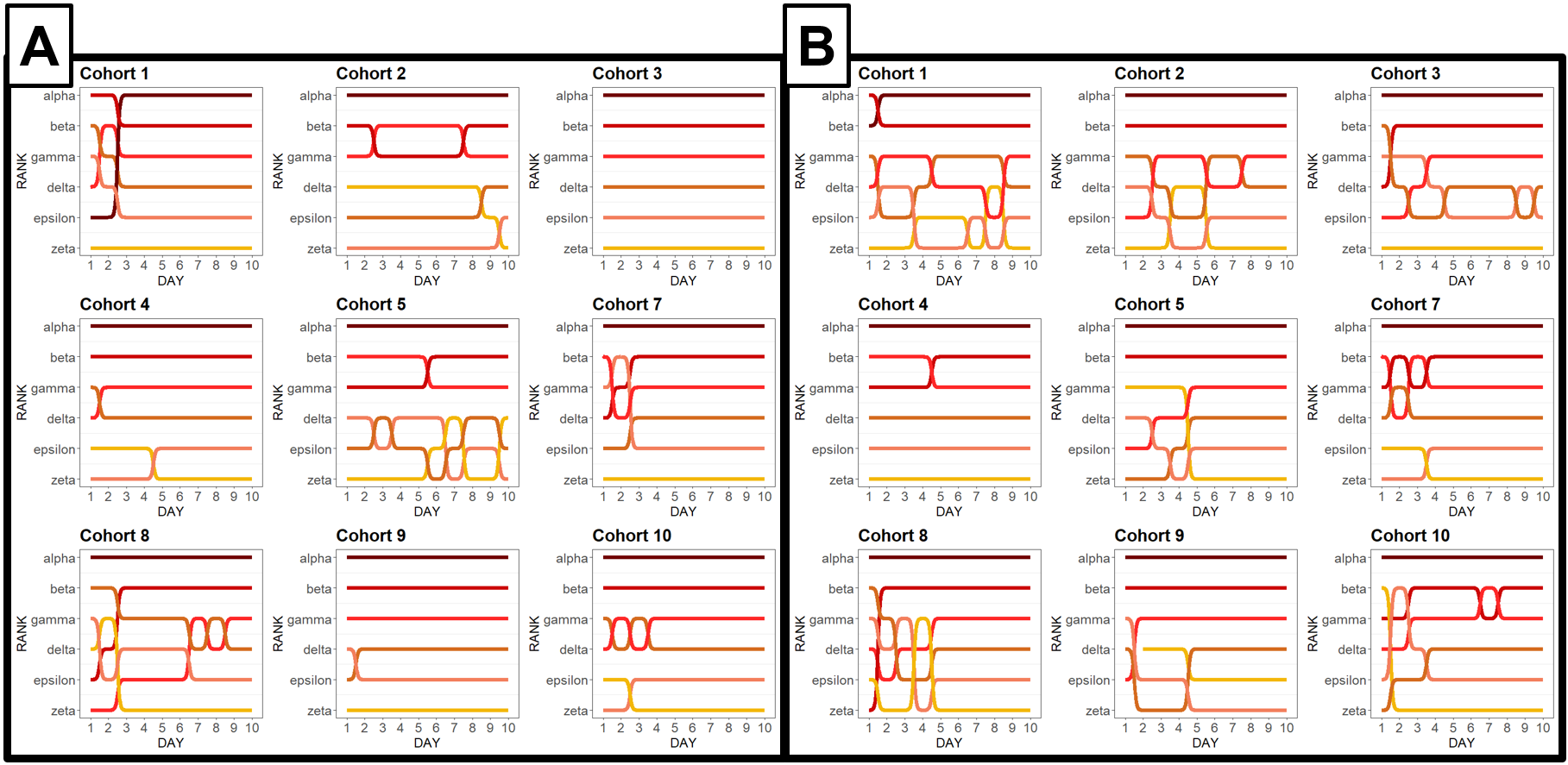

**Supplemental Figure 5: (A) Hierarchy rankings at the end of each day using observer coded agonistic interactions.** Rank progressions at the end of each day of 9 days of social observation for all males are presented. Colors represent the hierarchy ranking on the final day, for each plot, not individual id. **(B) Hierarchy rankings at the end of each day using automated follow tracking.** Rank progressions at the end of each day of 11 days of group housing for all males are presented. For 6/9 cohorts, the top three ranks of the cohort were established by Day 6 of group housing. For all cohorts, the top three ranks of the cohort were established by Day 9 of social group housing.

**Supplemental Figure 6**

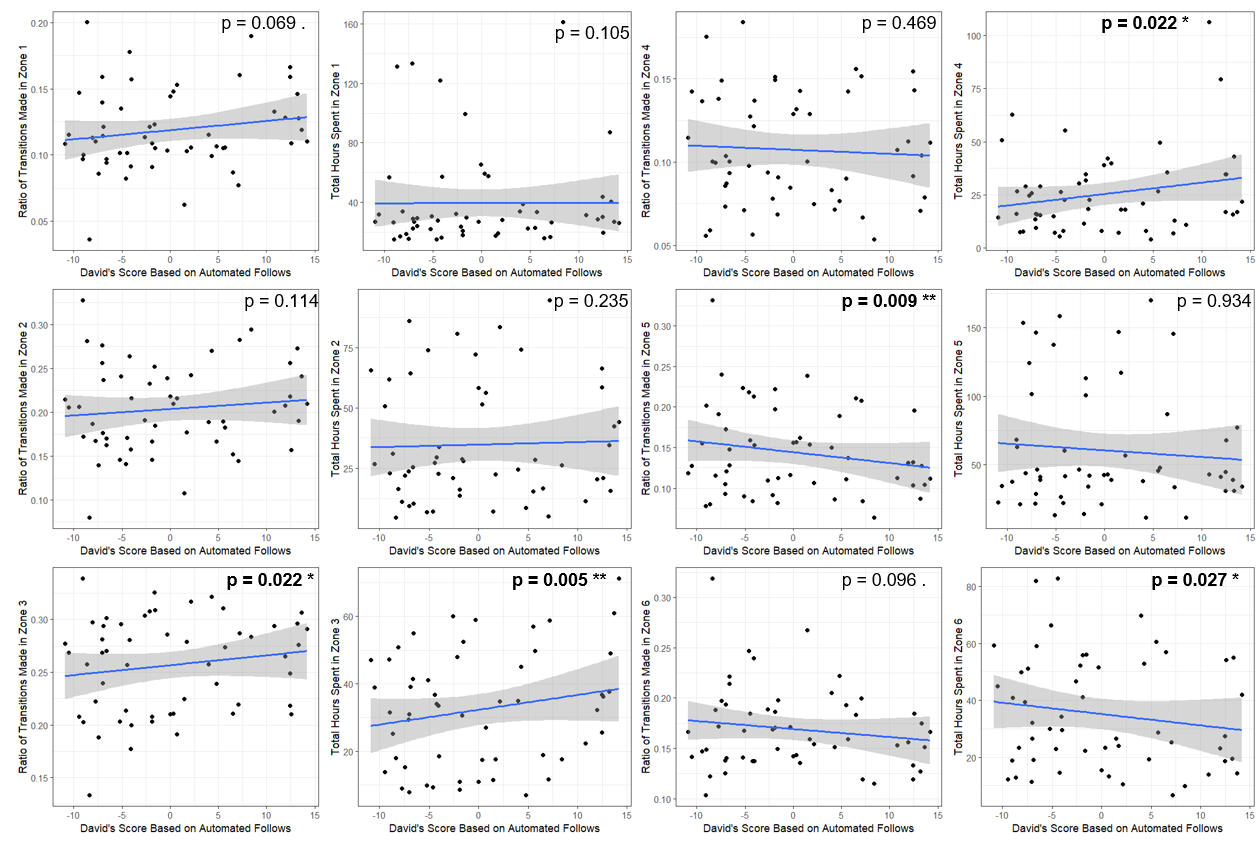

**Supplemental Figure 6: The relationship between dominance rating (David’s Scores from automated follow data) and (left) ratio of transitions in a specific zone per all transitions made, or (right) total hours spent in a specific zone.** Higher-ranking mice spent more time and made a greater proportion of transitions in zones 3-4, with a trend toward greater use of the food cage (zone 1), whereas lower-ranking mice showed greater use of the more peripheral zones 5-6.

**Supplemental Figure 7**

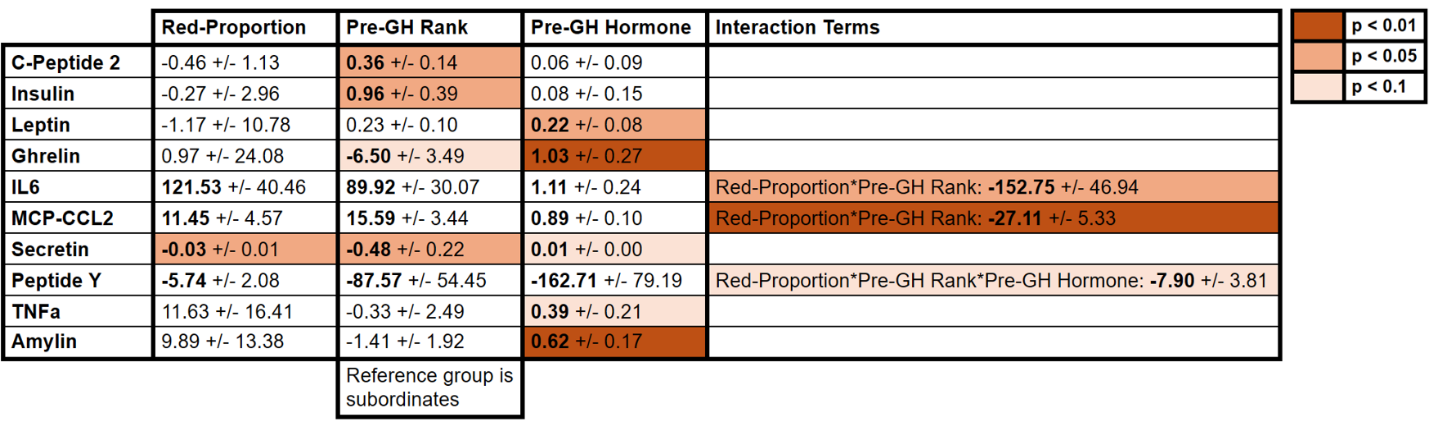

**Supplemental Figure 7: Estimates +/- Standard Errors from all post-group house concentration predicting models.** Presented above are the model estimates and standard errors for all hormones measured. Predictor variables include the proportion of tube transitions made in red light divided by the number of total tube transitions made in white and red light conditions (Red-Proportion), the hierarchical rank of mice prior to group housing when mice were housed in pairs (Pre-GH Rank), and the hormone concentrations of all hormones measured prior to group housing (Pre-GH Hormone). Interaction effects were tested for all predictors and significant interactions are recorded in the Interaction Terms column. The darkest red cells are predictor variables that demonstrated a p value of less than 0.01, the medium red cells are predictor variables that demonstrated a p value of less than 0.05 but more than 0.01. And the light red cells represent predictor variables that demonstrated a p value of less than 0.1 but more than 0.05. In models where interaction terms were significant, estimates are presented but they are warped by the presence of an interaction effect in the model so significance of those predictors is not presented.

**Supplemental Figure 8**

**A)**

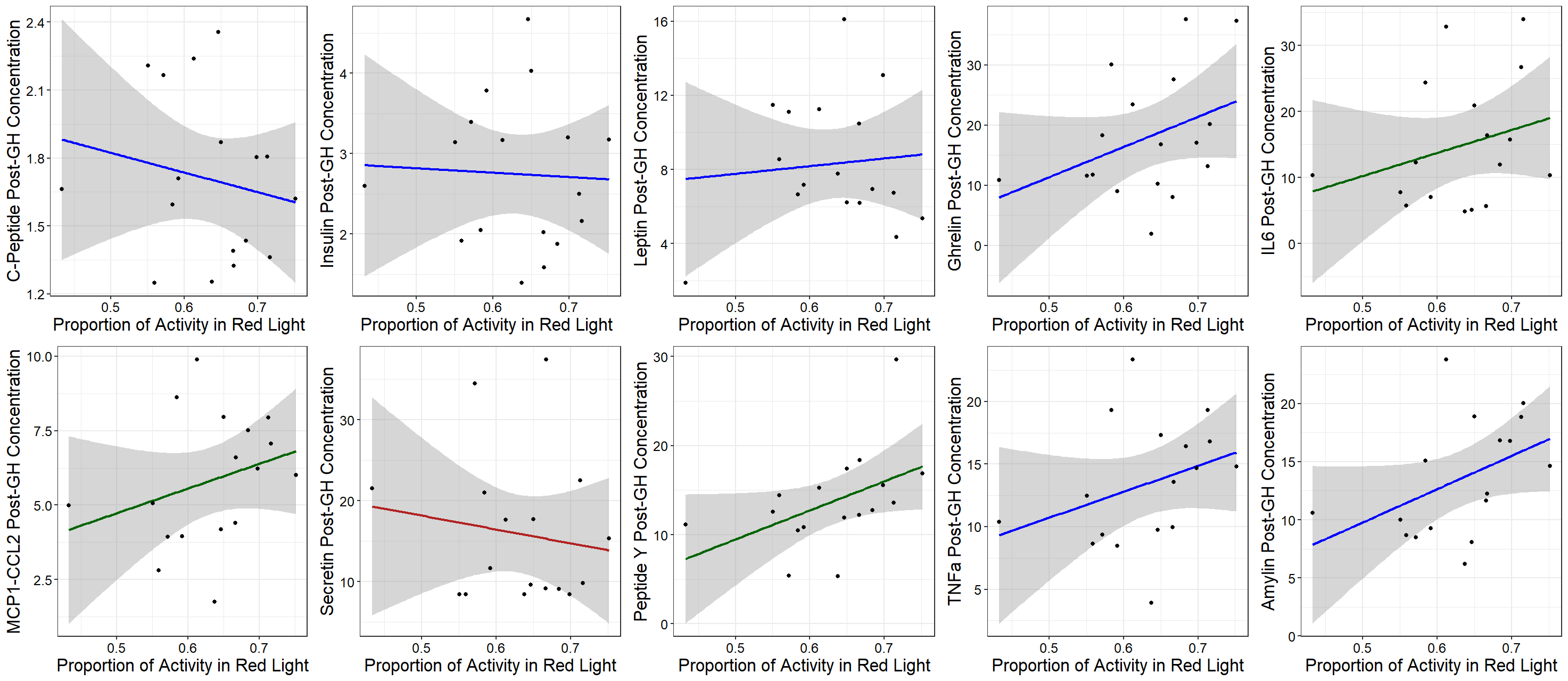

**B)**

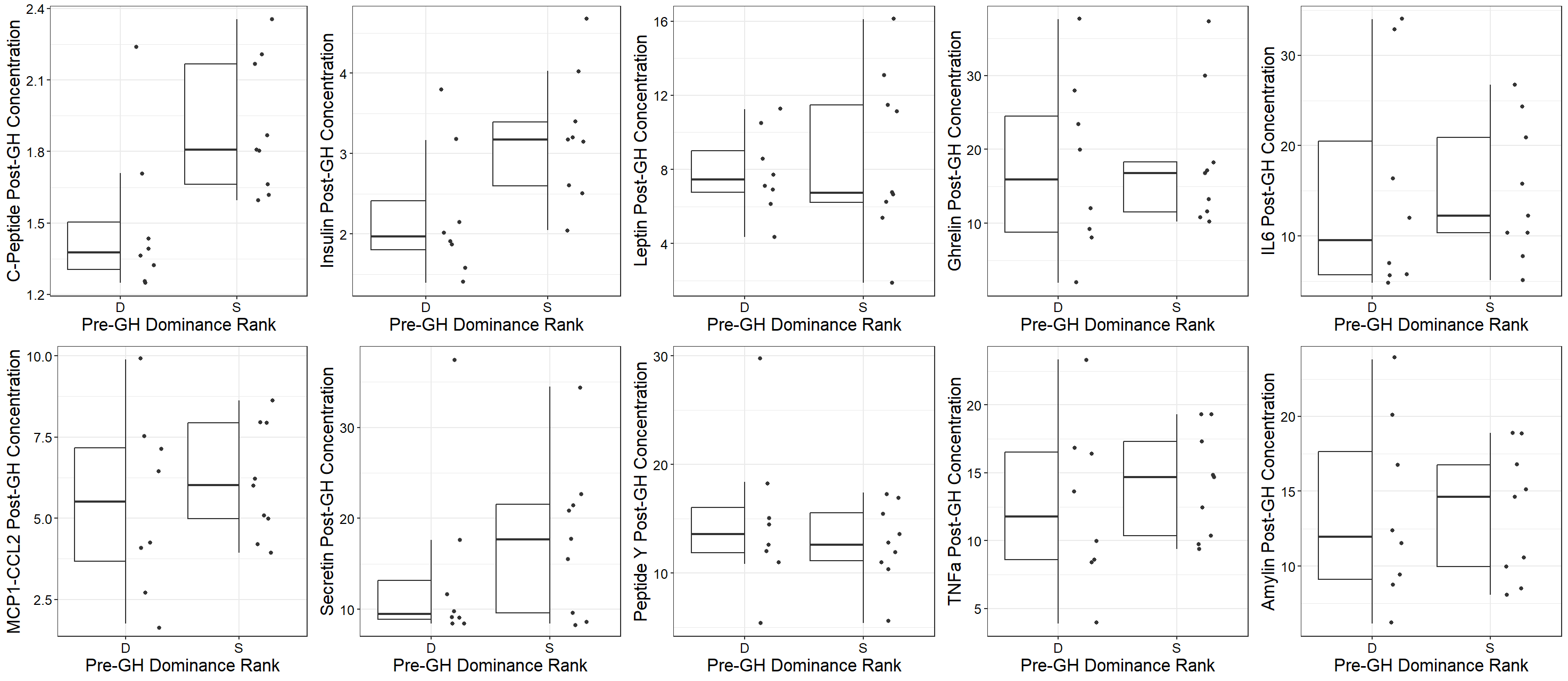

**Supplemental Figure 8: (A) The relationship between proportion of tube transitions in the red light compared vs all transitions made and hormone concentration after group housing. (B) The relationship between pre-group house ranking and hormone concentration after group housing.** A significant relationship between proportion of transitions in red light vs. all transitions (higher values indicating better circadian rhythmicity) was found in post group housing Secretin (indicated by dark red line). The proportion of activity in the red light is involved in a significant interaction with at pre-group housing dominance rank in IL6, MCP1-CCL2, and Peptide Y and also with pre-group housing Peptide Y hormone concentration in Peptide Y. As such, the statistical values of these predictors cannot be considered individually (indicated by dark green line). Other hormonal relationships with red light transition proportion are not significant. A significant relationship was observed between pre-group housing rank and C-Peptide, Insulin, and Secretin hormone concentration. A marginally significant relationship was identified between pre-group housing rank and post-group housing Ghrelin concentration, but the effect was small. An interaction between pre-group housing dominance rank and pre-group housing hormone concentration was observed in Peptide Y (see Supplemental Figure 7 for all model estimates, standard errors, and significance values).
